# A familial missense variant in the Alzheimer’s Disease gene *SORL1* impairs its maturation and endosomal sorting

**DOI:** 10.1101/2023.07.01.547348

**Authors:** Elnaz Fazeli, Daniel D. Child, Stephanie A. Bucks, Miki Stovarsky, Gabrielle Edwards, Shannon E. Rose, Chang-En Yu, Caitlin Latimer, Yu Kitago, Thomas Bird, Suman Jayadev, Olav M. Andersen, Jessica E. Young

## Abstract

The *SORL1* gene has recently emerged as a strong Alzheimer’s Disease (AD) risk gene. Over 500 different variants have been identified in the gene and the contribution of individual variants to AD development and progression is still largely unknown. Here, we describe a family consisting of 2 parents and 5 offspring. Both parents were affected with dementia and one had confirmed AD pathology with an age of onset >75 years. All offspring were affected with AD with ages at onset ranging from 53yrs-74yrs. DNA was available from the parent with confirmed AD and 5 offspring. We identified a coding variant, p.(Arg953Cys), in *SORL1* in 5 of 6 individuals affected by AD. Notably, variant carriers had severe AD pathology, and the *SORL1* variant segregated with TDP-43 pathology (LATE-NC). We further characterized this variant and show that this Arginine substitution occurs at a critical position in the YWTD-domain of the *SORL1* translation product, SORL1. Functional studies further show that the p.R953C variant leads to retention of the SORL1 protein in the endoplasmic reticulum which leads to decreased maturation and shedding of the receptor and prevents its normal endosomal trafficking. Together, our analysis suggests that p.R953C is a pathogenic variant of *SORL1* and sheds light on mechanisms of how missense *SORL1* variants may lead to AD.

## Introduction

Alzheimer Disease (AD) is the most common cause of dementia worldwide. The etiology of AD remains elusive, slowing development of disease modifying therapies. Pathogenic variants in *PSEN1*, *PSEN2* and *APP* are associated with autosomal dominantly inherited early-onset AD (ADAD), although those families are rare and make up only a very small fraction of all AD. Nevertheless, knowledge gained from studying ADAD has been valuable to our understanding of the clinical, pathological and mechanistic features of AD more broadly. Late onset AD also has a genetic component and is known to be highly heritable, estimated at 60-80%[33] and heritability can vary with age[9]. Genome wide association studies (GWAS) as well as genome and exome sequencing studies have revealed the complexity of biological processes contributing to AD risk and progression[70]. Given that families with AD likely harbor at least one AD genetic risk factor, they can provide important insight into genetic risk and disease pathogenesis.

The Sortilin-like receptor, *SORL1,* (protein: SORL1/SORLA) was originally identified as a member of the LDL receptor family, and the SORL1 protein is now classified as one of five mammalian sorting receptors called VPS10p receptors[31, 32, 72–74]. SORL1 functions as an endosomal receptor to assist cargo sorting out of the endosome to either the cell surface via the recycling pathway or to the trans-Golgi network (TGN) via the retrograde pathway[21, 24, 34, 51, 65]. For sorting of AD-related cargo, including Amyloid-β peptide (Aβ) and APP, SORL1 directly interacts with the multi-sorting complex retromer, itself highly implicated in endo-lysosomal health and neurodegeneration[18, 22, 57].

Through both candidate gene studies and GWAS, *SORL1* was found to be a strong genetic risk factor for AD[42, 43, 58, 59]. Exome-sequencing studies have shown that rare loss-of-function S*ORL1* alleles, leading to haploinsufficiency, have been associated with highly penetrant AD[25, 26, 55, 56, 71], although the full breadth and contribution of *SORL1* variants in AD is not fully defined. A large number (>500) of *SORL1* variants have been identified in patient populations with AD, but with variable levels of evidence for pathogenicity. Recently, two missense variants have been associated with autosomal dominant AD: p.(Asp1545Val) (Bjarnadottir et al., Manuscript in preparation) and p.(Tyr1816Cys)[35]. In case of the p.(Tyr1816Cys) we showed how this mutation has only minor impact on the intracellular localization per se, but strongly decreased receptor dimerization in endosomes and retromer-dependent recycling to the cell surface[35]. Reported *SORL1* variants span the length of the gene and functional domains, and how different pathogenic variants impair the overall functions of SORL1 as an endosomal sorting receptor is not yet clear. It has been suggested that SORL1 maturation, which is a distinct change in some of the *N*-glycans attached to the luminal SORL1 domain[14], is decreased for some *SORL1* missense variants[50, 60]. Defining the biochemical consequences of pathogenic *SORL1* missense variants can shed light on mechanisms of disease involving SORL1 and other components of the endo-lysosomal network (ELN).

We present here a family with early and late onset AD in two generations. Genetic testing confirmed a novel *SORL1* variant, c.2857C>T p.Arg953Cys (R953C; NM_003105.5) which affects a residue in one of the repeats in the YWTD-domain, in 6 out of 7 affected individuals tested. Neuropathological studies demonstrated severe AD pathology, including cerebellar amyloid plaques, cortical neurofibrillary tangles, and TDP-43 deposition despite a young age of onset in most carriers of the *SORL1* R953C variant. One individual, I-2, was affected with AD but did not carry the SORL1 variant and did not show TDP-43 deposition. To further characterize this genetic variant, we turned to a previously described disease-mutation domain-mapping approach that relies on identified pathogenic variants in homologous proteins including members of the LDLR family[8], to predict pathogenicity based on the domain position at which the variant occurs in SORL1. We next generated a plasmid containing the p.R953C variant and transfected it into HEK293 and N2a cells. Our *in vitro* studies suggest reduced SORL1 maturation and impaired endosomal localization, confirming a functional consequence of the missense variant. The influence of the variant on SORL1 cellular localization may lead to impairment of endosomal sorting and have pathogenic effects. This study adds to the growing body of literature supporting a role for *SORL1* variants that may contribute to the missing AD heritability.

## Methods

### Study Participants

The family was ascertained by the University of Washington Alzheimer Disease Research Center. The study was approved by the UW Institutional Review Board (IRB) and all participants provided written consents.

### Genetic Studies

Genetic analysis was performed by the Northwest Clinical Genomics Laboratory (NCGL), a CLIA certified laboratory at the University of Washington. Samples underwent next-generation exome sequencing and analysis. Libraries were constructed according to the NCGL protocol. The KAPA Hyper Prep DNA library kit (KAPA Biosystems, Wilmington, MA, USA) was used to prepare the libraries, which were subsequently enriched using an in-house, optimized xGen Exome Research Panel v1.0 (Integrated DNA Technologies, Coralville, IA, USA). Paired-end sequencing of the exome-enriched libraries was performed on a HiSeq 4000 instrument (Illumina, San Diego, CA, USA). Greater than 99% of the coding regions and canonical splice sites were sequenced to a read coverage of at least 20X or greater. The average mean depth of coverage was 144 reads. Resulting sequences were aligned to the human genome reference (hg19) using the Burrows-Wheeler Aligner (BWA)[46]. Variants were identified using the Genome Analysis Toolkit (GATK)[19, 49]and were annotated using the SnpEff annotation tool[15] in combination with various population databases and variant impact scoring tools. Individual II-5 was initially screened with a 39-gene dementia panel which included: *APP*, *ARSA, APOE, ATP13A2, CHCHD10, CHMP2B, CSF1R, DNMT1, EIF2B1, EIF2B2, EIF2B3, EIF2B4, EIF2B5, FUS, GALC, GRN, HEXA, ITM2B, LMNB1, MAPT, NOTCH3, NPC1, NPC2, OPA1, PDGFB, PDGFRB, PLP1, PRNP, PSEN1, PSEN2, SLC20A2, SLC25A12, SORL1, TARDBP, TBK1, TBP, TREM2, TYROBP, VCP* which identified the *SORL1* p.R953C variant. Whole exome sequencing was then performed on II-1, II-2, II-4 and II-5 to evaluate for any other candidate variants and to investigate which variants segregated with the phenotype. Shared variants were filtered based on population data and variants with an allele frequency greater than 0.001 in ExAC were excluded from further analysis. Variants were manually evaluated through literature searches in PubMed. II-1, II-2, II-4 were also found to carry the *SORL1* p.R953C variant via exome sequencing. No other variants associated with dementing disorders were identified. II-3 was later found to carry the variant using Sanger sequencing.

### APOE genotyping

APOE genotyping was performed as previously published[44]. Briefly, genomic DNA was amplified in a 9700 Gene Amp PCR System (Applied Biosystems) using primers that amplify *APOE* gene’s exon 4. This PCR amplicon includes both the codon 112 (ε2/ε3 vs. ε4) and codon 158 (ε2 vs. ε3/ε4) polymorphic sites.

#### Taqman assay

SNPs rs429358 (ε2/ε3 vs. ε4) and rs7412 (ε2 vs. ε3/ε4) were genotyped using assay C_3084793_20 and assay C_904973_10 (Thermo Fisher), respectively. All reactions were carried out in a 9700 Gene Amp PCR System with a profile of 50°C for 5 minutes; 95°C for 5 minutes; 50 cycles of 95°C for 15 seconds, and 60°C for 1 minute.

#### Sanger sequencing

The PCR reaction/amplicon (1 µl) was used in BigDye sequencing reaction (Thermo Fisher) with a final volume of 10 µl. All reactions were carried out in a 9700 Gene Amp PCR System with a profile of 94°C for 1 minute; 35 cycles of 94°C for 30 seconds, 55°C for 10 seconds, and 60°C for 4 minutes; and a final extension of 60°C for 5 minutes. The PCR generated sequencing products were further purified using EDTA/ethanol precipitation and then subjected to DNA sequencing run using SeqStudio (Thermo Fisher). The sequencing data (electropherograms) were transferred and uploaded onto the Sequencher program (Genecodes) for sequence alignment.

Primer sequences:

APOE_Ex4_F: 5’ TCGGAACTGGAGGAACAACT 3’

APOE_Ex4_R: 5’ GCTCGAACCAGCTCTTGAGG 3’

### SORL1 genotyping

*SORL1* variant genotyping was performed on I-2, II-2, II-3, and III-6. Genomic DNA was amplified with Phusion Flash (Thermo Fisher) on a C1000 Touch Thermo cycler (BioRad) using primers that amplify exon 20 in *SORL1.* Cycle conditions: 98°C for 10s; 98°C for 1s, 65°C for 5s, 72°C for 10s X25 cycles; 72°C for 1 min. Cleaned PCR reactions were sent for Sanger sequencing using GeneWiz (Azenta Life Sciences). Sequences were examined manually using 4 Peaks software.

Primer sequences:

*SORL1* F: 5’ GCCTGGGATTTATCGGAGCA 3’

*SORL1* R: 5’ TGGCATCCCTCCATAGGCT 3’

### Neuropathology

Consent for autopsy was obtained from the donor or from the legal next of kin, according to the protocols approved by the UW Institutional Review Board. At the time of autopsy, the brain was removed in the usual fashion. For patients I-2, II-2, II-3 and II-4, the left halves were coronally sectioned and samples were frozen for possible biochemical studies and the right halves were fixed in formalin. For patients II-1 and II-5, the entire brain was fixed in formalin. After fixation, the cerebrum was sectioned coronally, the brainstem was sectioned axially, and the cerebellum was sectioned sagittally.

Representative sections for histology were selected and evaluated according to National Institute of Aging-Alzheimer’s Association (NIA-AA) guidelines[52]. A microtome was used to cut 4 μm-thick tissue sections from formalin-fixed, paraffin-embedded tissue blocks. Hematoxylin and eosin (H&E), Luxol fast blue (LFB), and Bielschowsky silver-stained slides were prepared. Using previously optimized conditions, immunohistochemistry was performed using a Leica Bond III Fully Automated IHC and ISH Staining System (Leica Biosystems, Wetzlar, Germany). The sections were immunostained with mouse monoclonal antibody against paired helical filament tau (AT8, 1:1,000 dilution) (Pierce Technology, Waltham, MA), mouse monoclonal against β-amyloid (6E10, 1:5,000) (Covance, Princeton, NJ), rat monoclonal against phosphorylated TDP-43 (ser409/ser410, 1:1,000) (Millipore, Burlington, MA), and mouse monoclonal against α-synuclein (LB509, 1:500) (Invitrogen, Carlsbad, CA). Appropriate positive and negative controls were included with each antibody and each run.

### Site-directed Mutagenesis

The R953C variant was inserted in *SORL1* pcDNA3.1 and *SORL1*-GFP pcDNA3.1 using site directed mutagenesis kit (QuikChange #200521) according to manufacturers’ instruction with the following pair of primers: 5-gga tca cgt tca gtg gcc agc agt gct ctg tca ttc tgg aca acc tcc-3 and 5-gga ggt tgt cca gaa tga cag agc act gct ggc cac tga acg tga tcc-3.

### Cell transfection and western blotting

Approximately 5x10^5^ HEK293 and N2a cells were seeded on 6-well plates and transiently transfected with expression constructs for *SORL1*-WT or *SORL1*-R953C, using Fugene 6 Transfection Reagent kit (Promega) according to manufacturers’ instructions. 48 hours post transfection, cell medium was changed to serum free conditional medium and after 48 hours, cells and media were harvested. Cells were lysed using lysis buffer (Tris 20mM, EDTA 10mM, TritonX 1%, NP40 1%). Media samples (30ml) and lysate samples(20ug) were mixed with NuPAGE LDS sample buffer (Invitrogen, #2463558) supplemented with β-Mercaptoethanol (Sigma) and separated on SDS-PAGE using 4–12% NuPAGE Bis-Tris gels (Thermo). Proteins were then transferred to nitrocellulose membranes (Thermo) and incubated for 1h at room temperature in Blocking buffer (Tris-Base 0.25M, NaCl 2.5M, skimmed milk powder 2%, tween-20 2%). Next, membranes were incubated overnight at 4°C with LR11 antibody 1:1,000 (BDBiosciences # 612633) to detect SORL1 and Beta actin 1:2,000 (Sigma #A5441) as loading control, followed by three washes for 5 minutes in washing buffer (CaCl_2_ 0.2 mM, MgCl_2_ 0.1 mM, HEPES 1 mM, NaCl 14 mM, skimmed milk powder 0.2%, Tween 20 0.05%) and 1 hour incubation with HRP-conjugated secondary antibody (1:1,500, Dako, #P0260) for 1 hour at room temperature. Membranes were washed 5 times for 5 minutes, incubated with FEMTO detection reagent (Thermo #34095) and visualized by iBright1500 scanner. Quantification was performed by densitometric analysis in ImageJ and data were plotted in Graphpad Prism 9.5.0.

### Flow cytometry

Cell surface and total receptor level were analyzed by flow cytometry in live, transfected HEK293 and N2a cells. Briefly, HEK293 and N2a cells were transiently transfected with either *SORL1*-GFP-WT or *SORL1*-GFP-R953C plasmids. Twenty-four hours after transfection, cells were collected by trypsinization, pelleted, and resuspended in phosphate-buffered saline (PBS pH 7.4). After 15min incubation in blocking buffer (PBS pH 7.4,0.5% BSA), cells were immunostained at 4°C with rabbit anti-soluble-SORL1 primary antibody followed by washing two times with PBS pH 7.4 and 30min incubation with Alexa-flour 647 secondary antibody in the absence of detergent followed by 3 times washing and finally resuspension in FACS buffer (PBS pH 7.4, 2% FBS, 1% Glucose). Cells were analyzed by NovoCyte 3000 flow cytometer equipped with three lasers and 13 fluorescence detectors (Agilent, Santa Clara, CA). GFP and Alexa Flour 647 fluorophores were excited by the 488 and 640 nm lasers, respectively. Results were analyzed using FlowJo™ v10.8.1 Software (BD Life Sciences).

### Immunocytochemistry and Confocal Microscopy

Approximately 5x10^4^ HEK293 cells were seeded on poly-L-lysine coated glass coverslips and transfected with expression constructs for *SORL1*-WT or *SORL1*-R953C using Fugene 6 Transfection Reagent kit (Promega). 24h post-transfection, cells were fixed with PFA 4% for 10 minutes at room temperature, followed by a wash in PBS pH 7.4. Coverslips were washed twice in PBS with 0.1% Triton-X 100 (for intracellular staining) or only PBS (for membrane staining) and later blocked for 30 minutes at room temperature in blocking buffer (PBS, FBS 10%). Cells were then incubated overnight at 4°C with pAb_5387 (a polyclonal rabbit serum generated for the entire SORL1 ectodomain[31]) antibody alone or with an antibody against markers specific for each intracellular compartment (EEA1 for early endosomes, TFR for recycling endosomes, and Calnexin for ER). Next, cells were washed in PBS with or without Triton-X 0.1 % and incubated in Alexa Flour secondary antibodies (Invitrogen, 1:500) for 1 hour at room temperature. After washing once in PBS, cells were stained with Höechst (Abcam, 1:50,000) for 10 minutes at room temperature. The coverslips were then mounted on glass slides using DAKO fluorescence mounting medium (Agilent) and were imaged using Zeiss LSM800 confocal microscope. Colocalization was quantified using the JACOP plugin in ImageJ software and presented as Mander’s correlation coefficient. Graphing and statistical analysis of the data were performed with GraphPad Prism 9.5.0. Antibodies used were as follows: rabbit polyclonal anti-SORL1 (pAb_5387; Aarhus University) 1:300, mouse monoclonal anti-SORL1 (mAb_AG4; Aarhus University) 1:100, anti EEA1(#610457 BDBiosciences) 1:100, anti TFR 1:100(# A-11130 Invitrogen), anti-Calnexin (1:100) (#610523 BDBiosciences).

### Statistical analysis

The data are represented as the mean ± s.d. The ‘n’ numbers represent the number of biological replicates in each experiment, while for imaging studies ‘n’ represents the total number of cells analyzed. Data was analyzed using parametric two-tailed paired (WB analysis and flow cytometry) or unpaired (immunostaining) t-tests. A P-value of less than 0.05 is considered statistically significant. All statistical analysis was completed using GraphPad Prism 9.5.0 software.

## Data Availability

The authors confirm that the data supporting the findings of this study are available within the article and/or its supplementary material or available from a corresponding author upon reasonable request.

## Results

### Clinical Description

Three generations (**Figure 1**) of the study family are presented here. Clinical features are reported in **Table 1**. Both parents (I-1 and I-2) developed late onset dementia and I-1 also demonstrated parkinsonism and aggressive behavior. Of the 5 individuals in the II generation sibship, 4 were clinically diagnosed with AD, with a range of age of onset from 51 years to 73 years. II-3 was reported to carry a clinical diagnosis of dementia prior to death and had age of onset 74 yrs. II-4 and II-5, identical twins, developed early onset AD at age 57 years and 51 years, respectively. II-5 developed aphasia and apraxia in addition to memory loss. III-6, daughter of II-2, developed progressive spasticity at age 44. She has also developed evidence of executive dysfunction determined by neuropsychiatric evaluation at age 45 and again on repeat testing at age 46 without progression. She has not shown any lower motor neuron findings or any other neurological signs. MRI brain did not show atrophy or other abnormality (data not shown).

**Figure 1.**
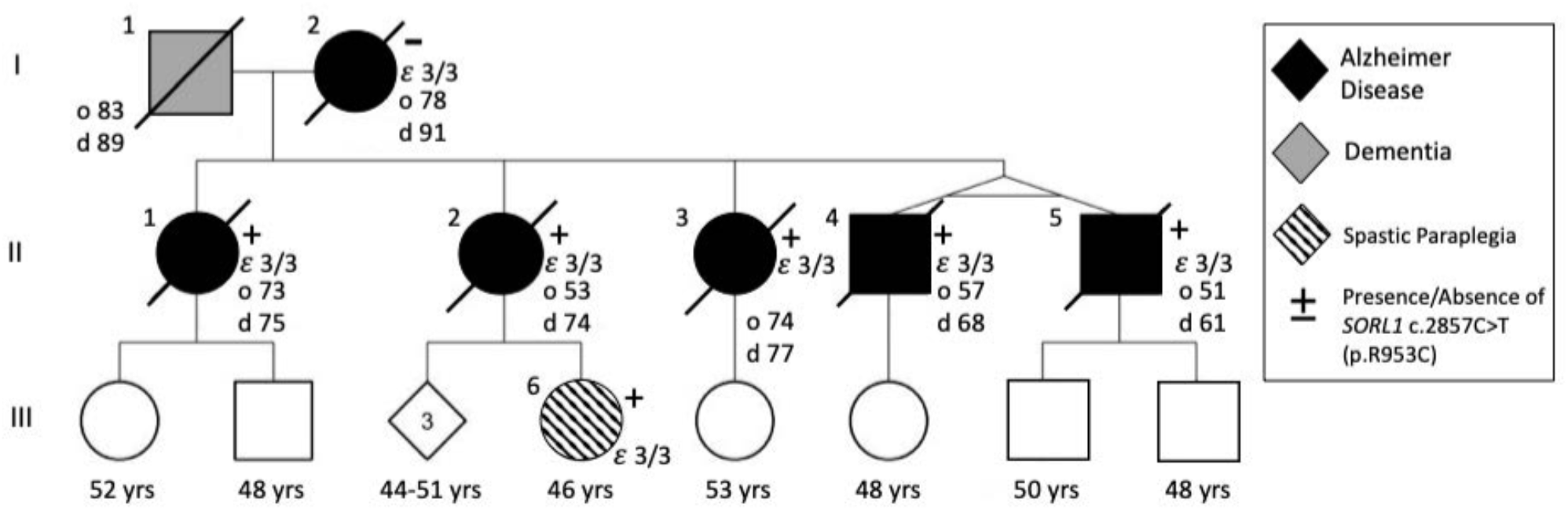
Pedigree of SORL1 R953C family: Solid black indicates individuals diagnosed with Alzheimer Disease which was confirmed by neuropathology. Dark grey indicates clinical diagnosis of dementia. Onset of disease (“o” years) and age at death “d” years is indicated next to the individual when applicable. Circles indicate female, square indicates male. Diamond is sex unknown to investigators at time of report. + or – indicates presence or absence of *SORL1* c.2857C>T variant. In individuals where APOE genotype was assessed it is indicated on pedigree.

**Table 1:**
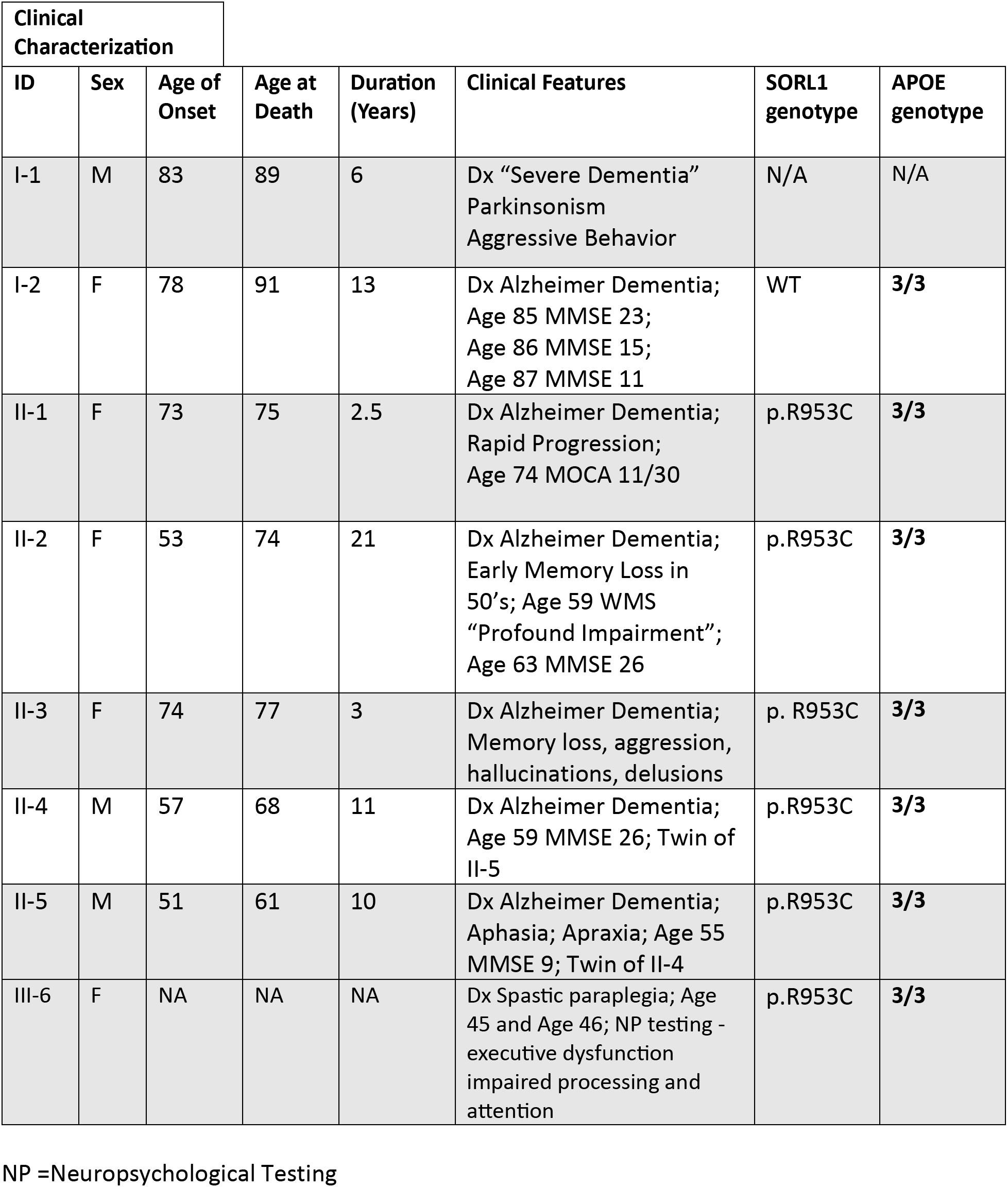
Clinical Characterization.

### Neuropathology

Individuals I-2, II-1, II-2, II-3, II-4, and II-5 were evaluated at autopsy, and findings are summarized in **Table 2**. Brain weight in all cases except II-1 was below the 10^th^ percentile for age and sex[10]. Atherosclerosis was present in all cases, with plaques extending beyond the first branch point of at least one cerebral artery (defined here as moderate); in case II-4, atherosclerotic plaques were also visible on the external surface and thus graded as severe. No other abnormalities were observed grossly in any case.

**Table 2:** Neuropathologic Findings.

| PEDIGREE<br>NUMBER | BRAIN<br>WEIGHT<br>(G) | ATHERO-SCLEROSIS | THAL<br>PHASE | BRAAK<br>STAGE | CERAD | ADNC | HIPPO-CAMPAL<br>SCLEROSIS | LATE<br>STAGE | LEWY<br>BODY<br>DISEASE |
| --- | --- | --- | --- | --- | --- | --- | --- | --- | --- |
| I-2 | 900 | Moderate | 4 | VI | Frequent | HIGH | Absent | 0 | Absent |
| II-1 | 1136 | Moderate | 4 | VI | Frequent | HIGH | Absent | 2 | Diffuse |
| II-2 | 867 | Moderate | 5 | VI | Frequent | HIGH | Present | 2 | Limbic |
| II-3 | 1098 | Moderate | 5 | V | Frequent | HIGH | Absent | 1 | Absent |
| II-4 | 950 | Severe | II-4 | VI | Frequent | HIGH | Present | 2 | Brain-<br>stem |
| II-5 | 1120 | Moderate | 5 | VI | Frequent | HIGH | Absent | 2 | Absent |

### Histopathology

All autopsy cases were evaluated by the standard NIA-AA protocol[30, 52]. β-amyloid plaques progressed to the midbrain in cases I-2 and II-1 (Thal phase 4 of 5), and extended to the cerebellum in cases II-2, II-3, II-4, and II-5 (Thal phase 5 of 5, **Figure 2a**). Tau tangles were present within the calcarine cortex/primary visual cortex in all cases (Braak and Braak stage VI of VI, **Figure 2b**). Cortical neuritic plaque density in all cases was frequent by Consortium to Establish a Registry for Alzheimer’s Disease (CERAD) criteria (**Figure 2c**). The features in each case meet criteria for high Alzheimer’s disease neuropathologic change (ADNC) by NIA-AA guidelines[52]. Additionally, all generation II cases had TDP-43 inclusions in the amygdala and hippocampus, consistent with limbic-predominant age-related TDP-43 encephalopathy neuropathologic change (LATE-NC) stage 1 or 2 out of 3[53] (**Figure 2d**); TDP-43 inclusions were not seen in case I-2 (*SORL1* variant negative). Hippocampal sclerosis was also seen in cases II-2 and II-4. Varying stages of Lewy body disease (LBD) were also identified, with diffuse (neocortical) LBD diagnosed in II-1, limbic (transitional) LBD in II-2, and brainstem-predominant LBD in II-4 (**Figure 2e**). We present neuropathological findings of all subjects that underwent brain autopsy in **Figure 3**.

**Figure 2.**
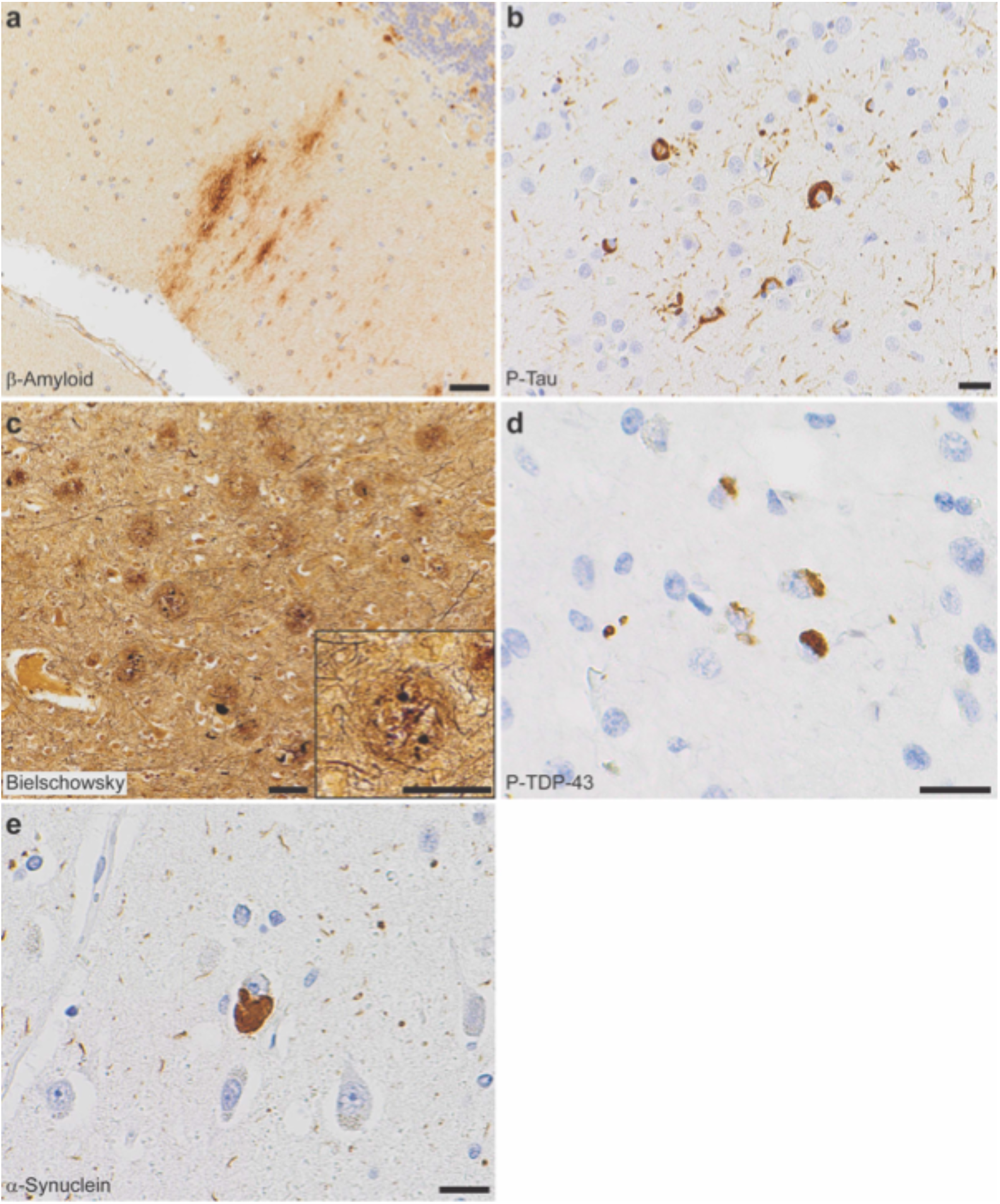
Neuropathologic evaluation demonstrates high Alzheimer disease pathologic change (ADNC) by NIA-AA criteria in SORL1 R953C cases. (a) Representative section of cerebellum stained for β-amyloid (6e10), highlighting plaques within the molecular layer and warranting a Thal phase 5. Patient II-5, scale bar = 50 µm. (b) Representative section of calcarine cortex stained for phosphorylated tau (P-Tau; AT8), highlighting neurofibrillary tangles in a background of dystrophic neurites, consistent with Braak and Braak stage VI. Patient II-5, scale bar = 20 µm. (c) Representative section of middle frontal gyrus stained with Bielschowsky silver demonstrating frequent neuritic plaques by CERAD criteria. Insert shows a representative neuritic plaque, composed of brown, targetoid β-amyloid associated with black dystrophic neurites. Patient II-5, scale bars = 50 µm. (d) Representative section of hippocampus stained for phosphorylated TDP-43 (P-TDP43), demonstrating intracytoplasmic inclusions and scattered dystrophic neurites. The pattern is consistent with limbic-predominant age-related TDP-43 encephalopathy (LATE) stage 2, though age < 80 years is atypical for sporadic LATE. Patient II-4, scale bar = 20 µm. (e) Representative section of anterior cingulate gyrus stained for α-synuclein, highlighting the presence of a Lewy body in a background of positive neurites. Though Lewy body disease was present in the majority of SORL1 R953C carriers, the pattern was highly variable. Patient II-2, scale bar = 20 µm.

**Figure 3.**
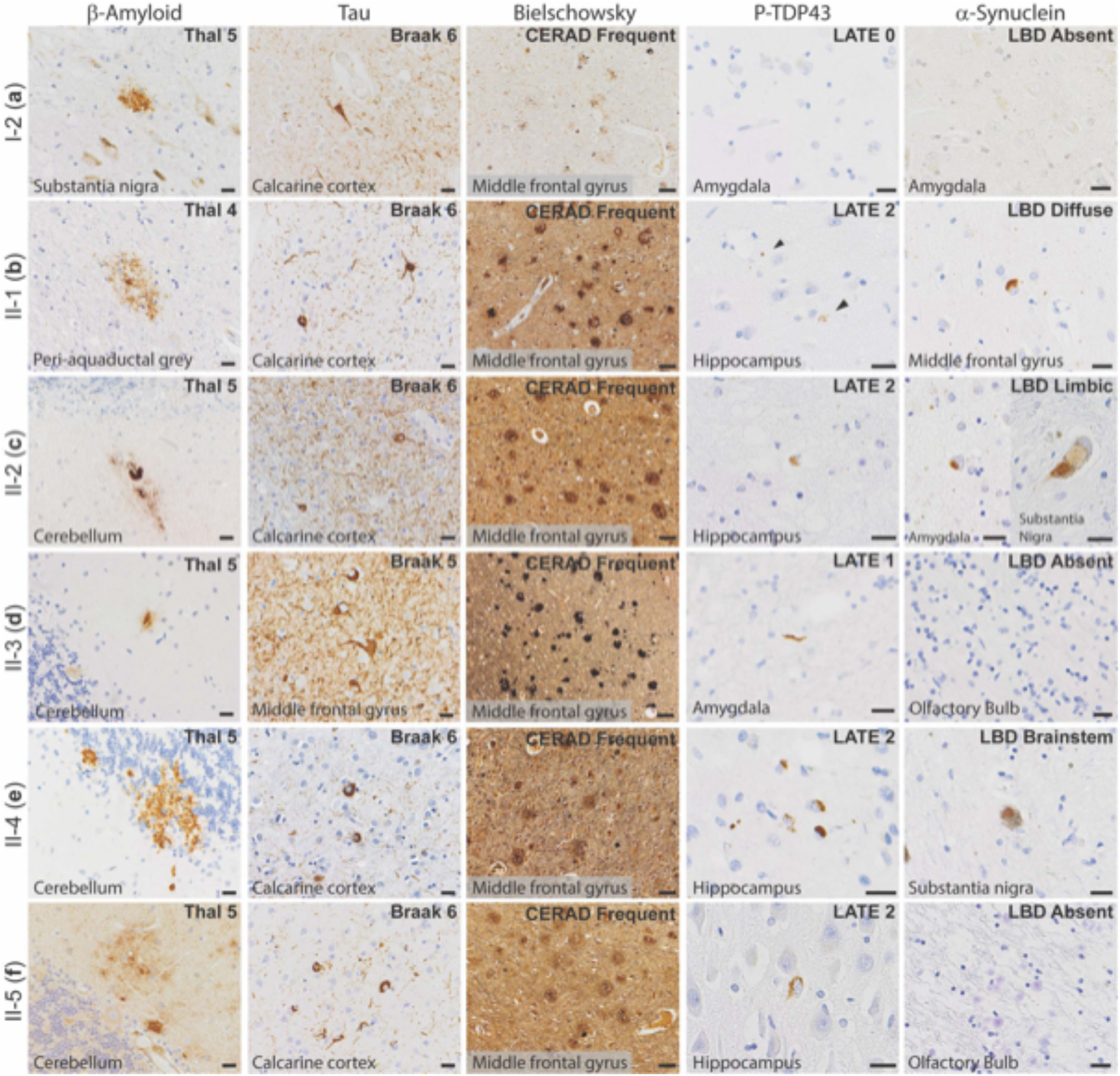
Neuropathology of all family members who consented to autopsy. Representative photomicrographs demonstrating highest level neuropathologic change in each autopsy case for β-amyloid plaques (6e10 antibody), neurofibrillary tangles (tau antibodies as listed below), neuritic plaques (Bielschowsky silver stain), phosphorylated-TDP-43 inclusions (P-TDP43 antibody), and Lewy bodies (α-Synuclein antibody). (a) Patient I-2, with β-amyloid plaques in the substantia nigra, neurofibrillary tangles (Tau2 antibody) in the calcarine cortex (primary visual cortex), and frequent neuritic plaque density by CERAD criteria (note that silver staining was lighter than other cases). No p-TDP43 or α-synuclein was present, shown here as lack of staining in areas affected early in disease process. (b) Patient II-1, with β-amyloid plaques in the periaqueductal grey matter of the midbrain, neurofibrillary tangles (AT8 antibody) in the calcarine cortex, and frequent neuritic plaque density by CERAD criteria. P-TDP43 inclusions were present in the hippocampus, highlighted by arrows. Lewy bodies were present in brainstem, amygdala, limbic structures, and frontal cortex (shown here). (c) Patient II-2, with β-amyloid plaques in the cerebellum, neurofibrillary tangles in the calcarine cortex (AT8 antibody), and frequent neuritic plaque density by CERAD criteria. P-TDP43 inclusions were present in the hippocampus. Lewy bodies were present in the amygdala and substantia nigra, consistent with a limbic (transitional) pattern. (d) Patient II-3, with β-amyloid plaques in the cerebellum, neurofibrillary tangles in the middle frontal gyrus (AT8 antibody), and frequent neuritic plaque density by CERAD criteria. P-TDP43 inclusions were present in amygdala neurites. No Lewy bodies were observed, demonstrated here by negative staining of the olfactory bulb, one of the earliest anatomic sites of Lewy body formation. (e) Patient II-4, with β-amyloid plaques in the cerebellum, neurofibrillary tangles in the calcarine cortex (Tau2 antibody), and frequent neuritic plaque density by CERAD criteria. P-TDP43 inclusions were present in the hippocampus. Lewy bodies were present in the pigmented cells of the substantia nigra but not in any other site. (f) Patient II-5, with β-amyloid plaques in the cerebellum, neurofibrillary tangles in the calcarine cortex (Tau2 antibody), and frequent neuritic plaque density by CERAD criteria. P-TDP43 inclusions were present in the hippocampus. No Lewy bodies were observed, again demonstrated here by negative staining of the olfactory bulb. Scale bars = 20 µm for β-amyloid, p-tau, p-TDP43, and α-Synuclein; scale bars = 50 µm for Bielschowsky silver stain.

### Genetic Findings

Due to early onset and family history of AD, subject II-5 underwent *PSEN1* and *APP* research genetic testing, which was negative in both genes. Years after the subject’s passing, his genetic material was included in an early onset AD cohort evaluated by an exome panel of 39 neurodegeneration genes. II-5 was found to carry a *SORL1* missense variant: NM_003105.5 c.2857C>T p.Arg953Cys (R953C). The reported allele frequency of this variant in gnomAD for those of European (non-Finnish) ancestry is 1/113646. It has not been reported in other populations assessed. In silico predictions varied; Polyphen: probably damaging, SIFT: tolerated, REVEL: 0.805, CADD v1.3: 25.4, PrimateAI: 0.633. No other pathogenic or likely pathogenic variants were identified in the other 38 genes on the neurodegeneration panel. Next, we screened II-1, II-2, II-4, and II-5 by whole exome sequencing, which revealed that all four subjects carry the *SORL1* R953C variant and no other pathogenic variants known to be associated with dementia were identified. We re-confirmed the presence of the *SORL1* R953C in II-2 variant using Sanger sequencing. II-3 passed away during preparation of this manuscript. We performed Sanger sequencing and confirmed the presence of the *SORL1* R953C variant in II-3. Using Sanger sequencing, we found that I-2 did not carry the *SORL1* variant, and no DNA samples were available from I-1. III-6 was found to carry the *SORL1* variant using Sanger sequencing of dermal fibroblasts. All Sanger sequencing results are presented in **Supplemental Figure 1**. C9orf72 gene expansion testing was negative in generation II and III-6. I-2, all individuals in the II generation and III-6 have an APOE ε3/ε3 genotype.

### Variant characterization

The arginine residue Arg953 is located at blade position 38 of the YWTD-domain repeated sequence, located within the fifth of six repeats that build the 6-bladed β-propeller domain of SORL1 (**Figure 4a**). We previously undertook a detailed disease-mutation domain-mapping approach to identify the most pathogenic sequence positions for the SORL1 domains and their risk for developing AD[8]. From this analysis, YWTD-domain sequence position 38 was identified as a high-risk site when arginine substitution occurs, and we identified variant p.Arg953His, (p.R953H) in three early-onset AD patients corresponding to the same SORL1 amino acid. However, the p.R953C variant was not identified in this large exome-sequencing study[25]. From previous disease mapping work[8], we identified 5 pathogenic variants in homologous proteins corresponding to substitution of an arginine at the YWTD-domain sequence position 38, summarized in **Table 3**. We report variant classification by VarSome, a search engine that aggregates databases, including ClinVar, and annotates pathogenicity of variants using the ACMG/AMP guidelines[41].

**Figure 4.**
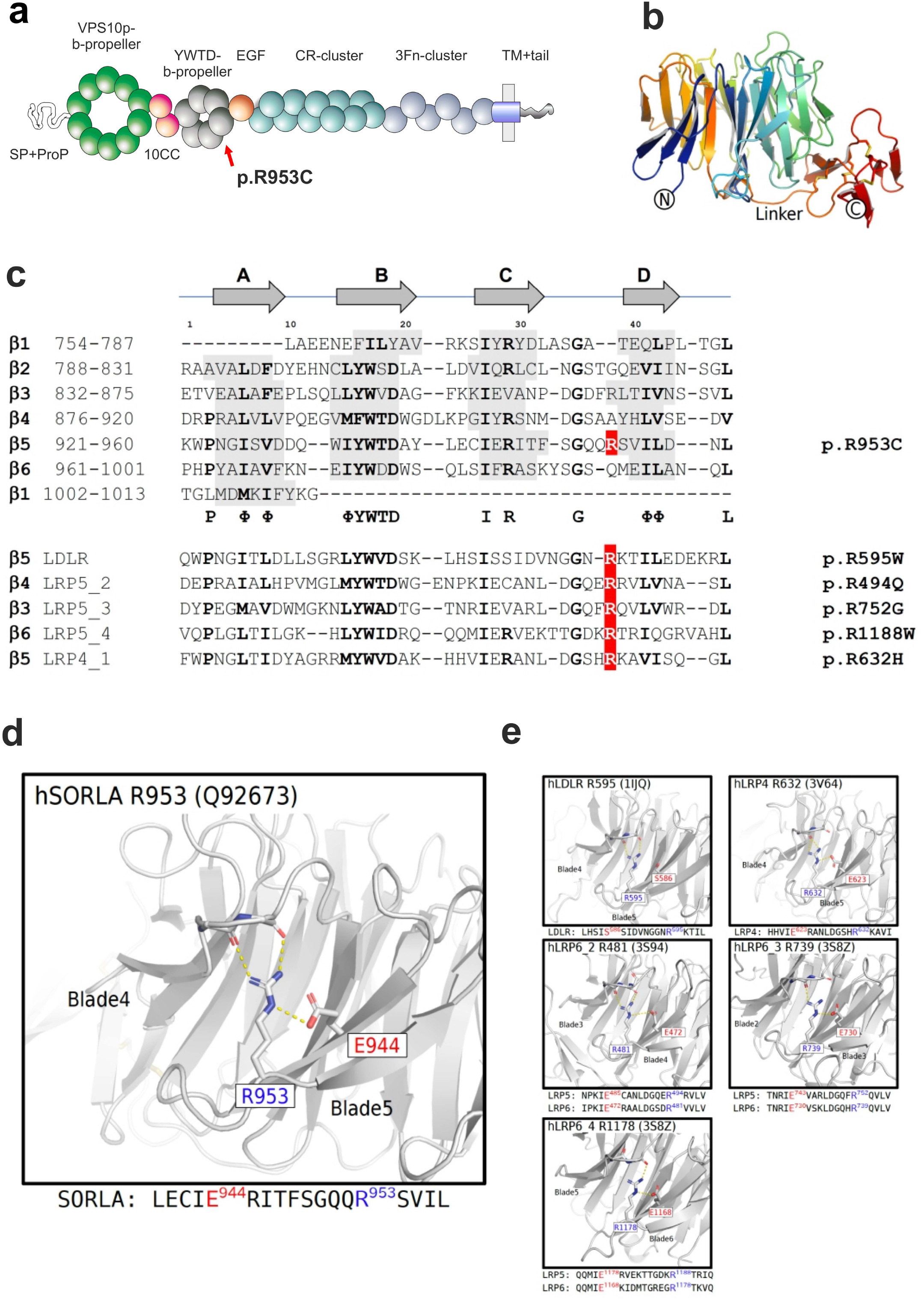
*In silico* characterization of SORL1 p. R953C. (a) Schematic presentation of the mosaic domain structure of the SORL1 protein comprising from the N-terminal end: VPS10p-domain with accompanying 10CCa/b domains, YWTD-repeated β-propeller domain (with p.R953C location indicated) with accompanying EGF-domain, cluster of 11 CR-domains, cluster of 6 3Fn-domains, a transmembrane domain followed by a cytoplasmic tail at the C-terminal end. (b) Three-dimensional model of the SORL1 YWTD-domain folding prepared from coordinates from ModelArchive (Y.Kitago, O.M. Andersen, G.A. Petsko. ModelArchive: https://modelarchiveorg/doi/10.5452/ma-agbg4). (c) Alignment of the ∼40 amino acids from each of the six YWTD-repeated sequences corresponding to the blades of the β-propeller with indication of β-strands in grey. The arginine R953 resides at domain position 38 of the sequence located in the loop between strands C and D of the fifth β-blade. Partly conserved domain positions are indicated with bold letters and the consensus residues below the SORL1 alignment. Below 5 sequences of YWTD-repeated sequences from homologous receptor proteins with known pathogenic variants corresponding to arginines at position 38. (d) The side chain of Arg-953 from SORL1 provides structural stabilization of the domain folding by an ionic interaction with the side chain of Glu-943 based on the three-dimensional model of the folded YWTD-domain. (e) Close-up of the Arg-Glu pairs from YWTD-domain crystal structures for residues in LDLR, LRP4 and LRP6 (LRP5 homolog) corresponding to pathogenic variants as listed in panel c

**Table 3:**
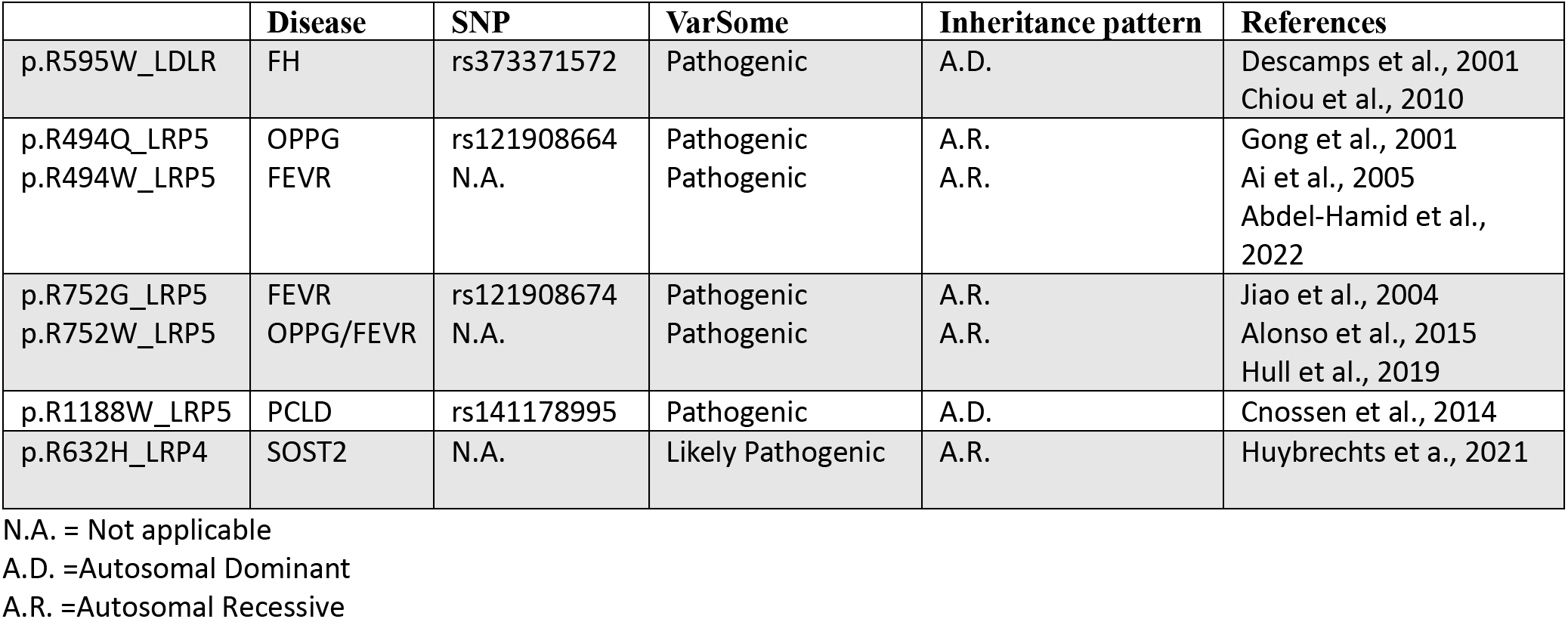
Homologous mutations for SORL1 R953C in LDLR and LRP.

Familial hypercholesterolemia (FH) is an autosomal dominant disorder with a prevalence of approximately 1 in 500 and most frequently caused by mutations in the gene for the low-density lipoprotein receptor (LDLR). The variant p.R595W^LDLR^ has been identified in patients with FH family history in cohorts from Belgium[20] and Taiwan[13] and considered an autosomal dominant variant.

Variants in another member of the LDLR gene family, LRP5, has been associated with a number of monogenic diseases, and different variants are often the cause of different clinical disorders. The p.R1188W^LRP5^ has been identified to segregate in a 40-member Dutch family with three generations of early-and late-onset cystogenesis inherited in an autosomal dominant fashion with Polycystic Liver Disease (PCLD). Cell-based studies to assess receptor activity confirmed significantly decreased activity of the mutated receptor compared to wild-type LRP5[16].

Osteoporosis-Pseudoglioma Syndrome (OPPG) is an autosomal recessive disorder and is caused by homozygous pathogenic variants in LRP5, due to the receptor function as a key regulator of bone metabolism through the Wnt signaling pathway. The biallelic presence of the pathogenic variant p.R494Q^LRP5^ has been identified as the cause of OPPG in families with homozygous carriers of the mutation[1, 3, 23]. Moreover, the p.R494W^LRP5^ that affects the same amino acid of LRP5 was identified as a potential pathogenic variant in a patient with Familial Exudative Vitreoretinopathy (FEVR), adding further support to the critical role of this amino acid to produce functional LRP5[47].

The variant p.R752G^LRP5^ was also identified as the cause of disease in a compound homozygous carrier for the FEVR autosomal recessive disorder [36]. Another variant that affects the same amino acid in LRP5; p.R752W^LRP5^ has been reported to associate with low bone mineral density in a female heterozygous carrier, and in combination with another pathogenic LRP5 variant (p.W79R that affects a YWTD-motif residue) in her son causes a severe case of compound heterozygous OPPG[4]. Moreover, the p.R752W^LRP5^ was identified as a potential pathogenic variant in a patient with FEVR when it was identified in a compound heterozygous carrier together with the pathogenic p.C1305Y variants in LRP5[27]. These studies add further support to the critical role of the arginine amino acid at domain position 38 to produce functional LRP5. A variant, p.R632H^LRP4^, affects the homologous receptor LRP4. This variant is causal for sclerosteosis when present as heterozygous compound mutation together with another pathogenic variant in LRP4 (p.R1170Q). Cell based assays confirmed how both of these mutations in LRP4 reduced receptor activity, providing support of the important function of the arginine also within the YWTD-domain of LRP4[29]. We summarize these findings and literature in **Table 3**.

We recently prepared a three-dimensional model of the SORL1 ectodomain including its YWTD-domain using the AlphaFold2 algorithm[34]. Here, we used this model to investigate the functional role of the arginine side chain (**Figure 4b, d**). From this model it is observed that the positively charged amino group makes ionic contacts with the side chain of the glutamic acid residue at blade-sequence position 28 (E944 of SORL1) serving to position the long arginine side chain in place to make further hydrogen bonds to two backbone carbonyls in the preceding loop between blades (**Figure 4d**), thereby strongly contributing to the folding and the stability of the entire six-bladed β-propeller domain. Interestingly, in four of the five blade-sequences containing the identified disease variants, a glutamic acid is similarly located at blade-sequence position 28 (**Figure 4c**).

Inspection of a larger alignment of YWTD-repeat sequences revealed that for most blade-sequences, a similar pattern is observed: when an arginine occupies blade-sequence at position 38, then a glutamate resides at blade-sequence position 28[8], suggesting this pair of residues may generally be important for the folding of YWTD-domains.

The crystal structures of the YWTD-domains have previously been solved for LDLR[61] and LPR4[75] including R595^LDLR^ and R632^LRP4^, the homologous residues for R953^SORL1^, respectively. The structure of LRP5 has not been determined, but as the crystal structure of the highly homologous LRP6 has been solved[2, 11, 12], it allowed us to use these YWTD-domain structures to gain insight in the functional role of the arginine side chain for the arginines at blade-sequence position 38 as well as for the LRP5 residues (R494^LRP5^/R481^LRP6^; R752^LRP5^/R739^LRP6^; R1188^LRP5^/R1178^LRP6^) (**Figure 4e**). Indeed, we found that the arginine side chains in each of the domains are binding backbone carbonyls in the n-1 linker, and for 4 of the 5 structures a salt bridge to a glutamic acid (at domain position 28) assist in keeping the arginine properly positioned to make the main chain interactions to the n-1 linker residue (**Figure 4e**). This supports a disease mechanism where substitution of the arginine may lead to domain misfolding and destabilization in general, and importantly also for R953 of SORL1.

### R953C disrupts SORL1 maturation and ectodomain shedding from the cell surface

SORL1 protein is synthesized in the ER and goes through a complex cellular process of maturation during trafficking in the ER and out of the Golgi into the ELN compartments and to the cell surface. The mature SORL1 isoform has complex-type *N*-glycosylations, and we previously showed only mature *N*-glycosylated SORL1 is shed from the cell surface to produce a fragment called soluble SORL1 (sSORL1)[14], and therefore a decrease in sSORL1 is often a direct measure of the maturation process being decreased for folding-deficient SORL1 mutant protein. Mature SORL1 migrates more slowly by SDS-PAGE, thus mature and immature isoforms of cellular SORL1 can be clearly distinguished[63] (**Figure 5a**).

**Figure 5.**
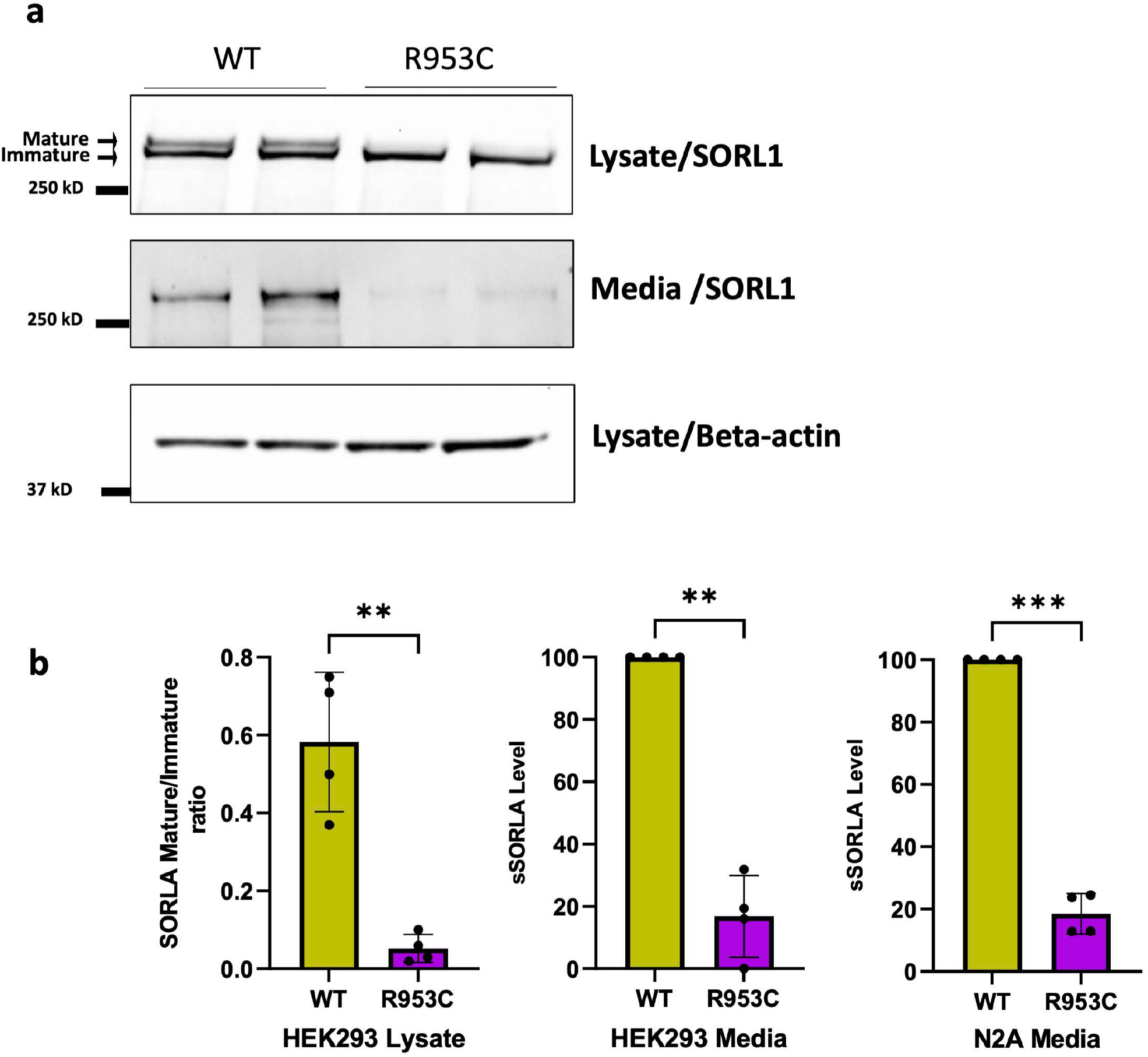
SORL1 R953C cells are defective in maturation and shedding of the SORL1 protein. (a) Representative western blotting of lysate and media samples from HEK293 cells transiently transfected with SORLA-WT or SORLA-R953C. (b) Densitometric analysis from HEK293 and N2a samples. The signal for the R953C is expressed relative to the WT signal. Results are expressed as Mean± SD and analyzed by parametric two-tailed paired t-test. Significance was defined as a value of **p<0. 01, ***p<0. 001. n= 4 independent experiments

To test whether the p.R953C variant affects SORL1 maturation and shedding, we transfected HEK293 and N2a cells with either the SORL1-WT or a SORL1-R953C construct. We performed Western blot analysis to determine the ratio of the mature to immature forms of the protein. We observed significantly decreased levels of mature SORL1 in HEK293 cells transfected with the R953C variant (**Figure 5a**). We next measured the level of sSORL1 in the culture medium of HEK293 and N2a cells, transiently transfected with expression constructs for SORL1-WT or SORL1-R953C. Compared to cells transfected with WT construct, we observed ∼ 80% reduction in the sSORL1 level in the media from both the tested cell types transfected with the R953C construct (**Figure 5a, b**)

### R953C reduces cell surface expression of SORL1

Because we observed a significant decrease in the shedding of SORL1-R953C, we tested whether the cell surface level of SORL1 could also be affected by this variant. We transiently transfected HEK293 cells with either SORL1-WT or SORL1-R953C and first analyzed cell surface levels of SORL1 using immunocytochemistry on unpermeabilized cells, which keeps the membrane intact to allow visualization of SORL1 protein solely located at the cell membrane. Using confocal microscopy, we observed considerably fewer cells expressing SORL1 at the cell surface in cells transfected with SORL1-R953C compared to SORL1-WT (**Figure 6a**). To quantitatively evaluate cell surface expression of SORL1-R953C relative to the total expression of the receptor in each individual cell, we used flow cytometry. We inserted the R953C variant into a C-terminally GFP tagged SORL1 construct, allowing for the detection of total expression of the receptor in each individual cell. We transfected both HEK293 cells and N2a cells and performed subsequent immunostaining of the transfected cells with anti-sSORL1 primary antibody and an Alexa Flour 647 secondary antibody in the absence of detergent to detect the cell surface expression of the receptor. These experiments demonstrated that more than 80% of the SORL1-R953C cells partially or completely retained SORL1 expression intracellularly compared to ∼10-15% of the SORL1-WT cells. Results were consistent in both HEK293 and N2a cells (**Figure 6b-c**).

**Figure 6.**
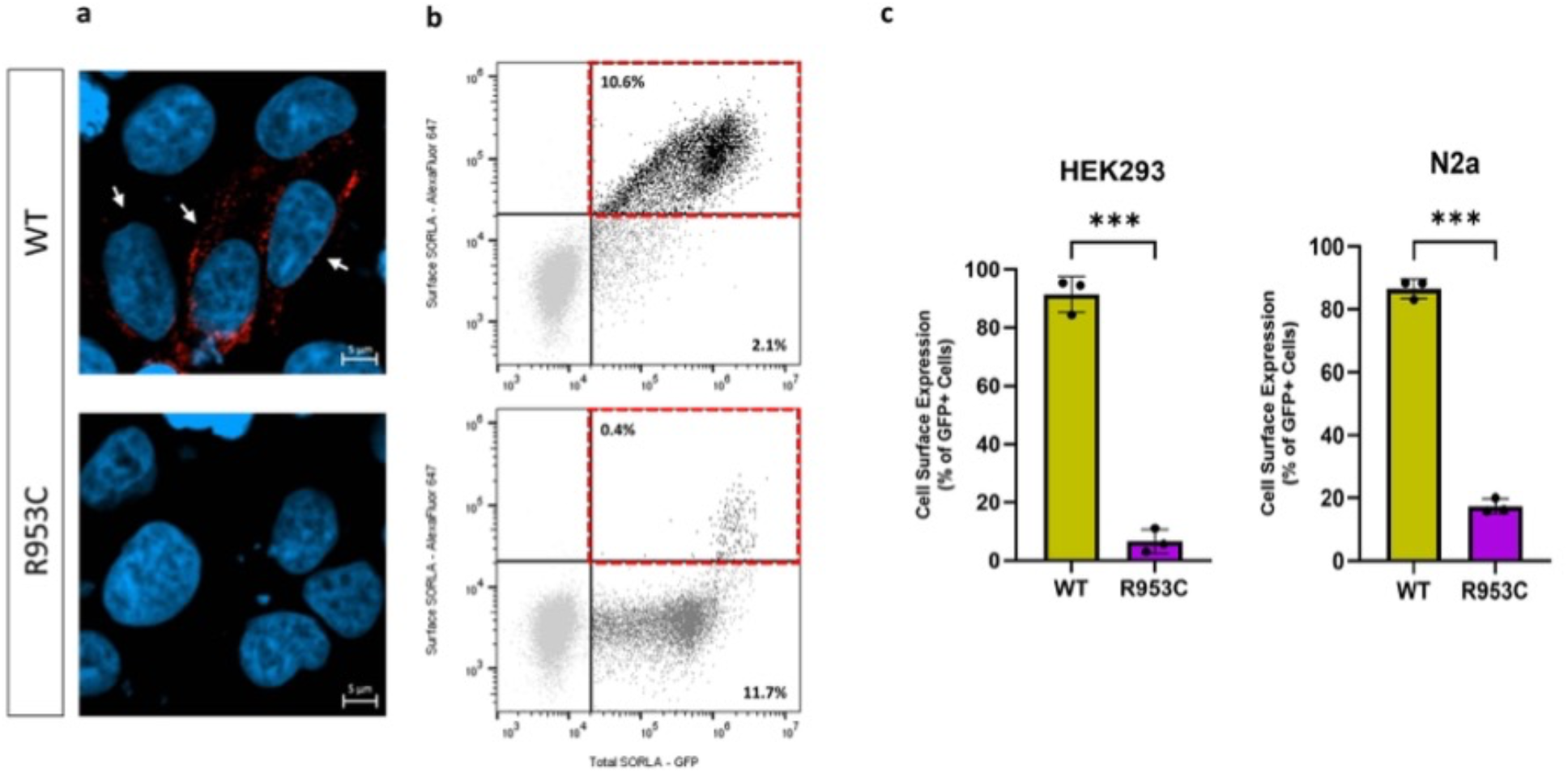
SORL1 R953C cells have reduced SORL1 protein localization on the cell surface. (a) Representative immunocytochemistry from HEK293 cells transiently transfected with SORLA-WT or SORLA-R953C expression construct and stained for SORLA (red) at the cell surface. White arrows show positive cells. (b) Flow cytometry dot plot showing surface (AlexaFluor 647 fluorescence) and total (GFP fluorescence) in live single HEK293 cells expressing WT-GFP and R953C-GFP. Vertical and horizontal lines represent thresholds for GFP and AlexaFluor 647-positive cells, respectively. Represented are GFP-positive cells with AlexaFluor 647 signal above (black, inside red dashed gate) or below threshold (dark grey); and untransfected cells (light grey). Numbers in the plots represent the percentages of the cells inside the gates. (c) Bar plots of AlexaFluor 647 fluorescence in HEK293 and N2a cells expressing WT-GFP or R953C-GFP, generated from population of GFP-positive cells. n=3 independent experiments. Results are expressed as Mean± SD and analyzed by parametric two-tailed paired t-test. Significance was defined as a value of ***p<0. 001.

### R953C prevents SORL1 from entering the endosomal recycling pathway

The differential cell surface localization and shedding of the R953C variant compared to WT led us to next investigate for possible changes in the intracellular localization of SORL1. For these experiments we transiently transfected HEK293 cells with either *SORL1*-WT or *SORL1*-R953C constructs. We analyzed co-localization of WT and R953C with two well-established endosomal markers, EEA1 (early endosome marker) and TFR (recycling endosome marker) and the ER marker Calnexin, 24 hours post-transfection. Using confocal microscopy, we demonstrated that the colocalization of R953C is strongly reduced with both endosomal markers (**Figure 7a-b**) and significantly increased in the ER (**Figure 7c**). Taken together, these data suggest that the R953C variant severely disrupts the normal cellular localization trafficking of SORL1 as would be expected if the mutation leads to defective protein folding.

**Figure 7.**
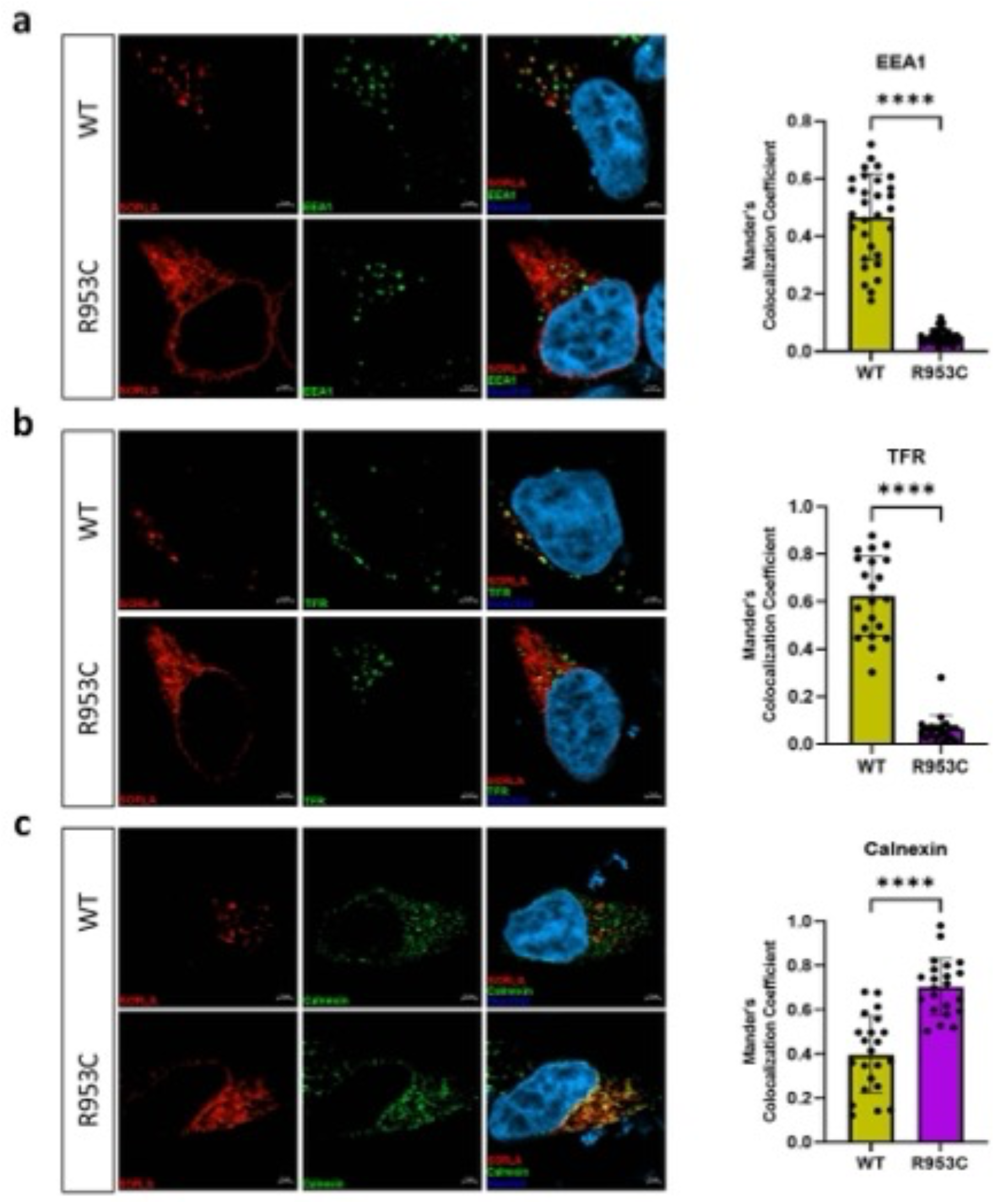
SORL1 R953C cells have reduced localization of the SORL1 protein in early and recycling endosomes. HEK293 cells transiently expressing WT or R953C (red) are shown for their colocalization with (a) EEA1 (early endosomal marker), (b) TFR (recycling endosomal marker), and (c) Calnexin (ER marker) (green). The nuclei were visualized with Hoechst (blue). Bar graphs on the right panel illustrate quantifications of colocalization between WT and R953C in cells co-stained for (a) EEA1, (b) TFR and (c) Calnexin. In all cases, the quantification of colocalization was represented as Mander’s correlation coefficient. 20-30 images per condition were analyzed. Data are shown as mean±SD and analyzed by parametric two-tailed unpaired t-test. Significance was defined as a value of ****p<0.0001.

## Discussion

*SORL1* is widely recognized as a strong AD risk gene though less is known about the AD risk attributable to rare missense variants[26, 55, 62]. Here we describe a family with two generations of both early and late onset AD in which we obtained brain autopsy pathology on 6 affected family members which enabled correlating clinical phenotype, genotype and neuropathology. Here we provide clinicopathological, genetic, and functional data supporting pathogenicity of a novel rare *SORL1* missense variant, p.(Arg953Cys) (R953C). Five of the five offspring were found to have the *SORL1* variant, and their age of onset ranged from 51yrs – 74 years. Notably, all affected individuals, including the mother who was WT for *SORL1* R953C, were of APOE3/3 genotype suggesting that APOE status was not contributing to risk or age of onset. Tissue from the mother (I-2) was analyzed by Sanger sequencing and found not to harbor R953C and tissue from the father was not available to confirm whether the allele was paternally inherited. Given the range of onset it is possible that additional genetic factors inherited from either parent has influenced expression of AD in both generations. Of note one living member of the family has been genotyped and is found to carry the variant (III-6) but is younger than the range of age of onset for the family. It is unknown whether her 3-year course of spasticity is related to the *SORL1* variant or is an unrelated case of a neurologic disease.

Neuropathology examination shows the presence of severe AD pathology, including extensive plaque and neurofibrillary tangle distribution. These histologic features typically correlate with advanced clinical disease[30, 52]. There are very few studies of neuropathology on *SORL1* variant carriers. There is one report of a *SORL1* homozygous truncating variant (c.364C>T, p.R122*) that shows severe cerebral amyloid angiopathy in addition to AD neuropathology as well as a patient with a splicing variant (c.4519+5G>A) in which AD was confirmed by neuropathological studies[5]. Yet another study shows SORL1 immunoreactivity in glial cells and white matter in a family with a *SORL1* variant c.3907C>T, p. R1303C.[69].

In our study, all cases underwent an extensive neuropathology examination in accordance with the most up-to-date guidelines for AD and related dementias[52, 53]. In this way, we were able to identify the presence of LATE-NC, marked by accumulation of TDP-43[53]. Interestingly, *SORL1* R953C segregated with LATE-NC pathology in 5 out of 5 offspring and with earlier age at AD onset in 3 out of 5 offspring. In fact, a recent analysis linked carrying a variant in *SORL1* with LATE-NC[38]. Although LATE-NC is a common co-pathology identified in AD, the underlying etiology of this TDP-43 pathology is not well understood. Age seems to be the strongest risk factor and it is most frequently noticed in individuals older than 80 years[53, 54]. Similar to other age-related neuropathologic changes, LATE-NC frequently co-occurs with other pathologies such as AD and/or hippocampal sclerosis[6], and its presence may accelerate the cognitive decline associated with these disorders[37]. It is worth acknowledging that it is possible that the co-morbid pathology of LATE-NC is driving the earlier age of onset in this family. Additionally, Lewy body disease (LBD) was frequently observed in *SORL1* R953C carriers (**Figure 3**). One other report in the literature has associated SNPs in *SORL1* with LBD, but also implicates SNPs in *APOE* and *BIN1* in this association as well[17]. While LBD limited to the amygdala is frequently observed in association with advanced ADNC[30, 52], ADAD due to *PSEN1*, *PSEN2* and *APP* have been associated with brainstem, limbic and diffuse LBD[45, 48] similar to what we find in this family. However, it is possible SORL1 itself contributes directly to synuclein pathology. Together, these co-pathologies suggest that SORL1 R953C may be mechanistically linked to multiple proteinopathies, clinically manifesting as AD but also impacting TDP-43[38] and α-synuclein histopathology[17]. While SORL1 and a-synuclein have not been shown to directly interact, a-synuclein is internalized via clathrin-mediated endocytosis and is present in many arms of the endo-lysosomal network[67]. Loss of TDP-43 function affects recycling endosomes and impairs trophic signaling in neurons[64]. Therefore, while there might not be a direct interaction, dysfunction of SORL1 as an endosomal receptor that facilitates endosome sorting pathways may lead to more global impairments in endo-lysosomal network function that affect other proteins involved in neurodegeneration.

We performed analyses to examine cellular and extracellular levels of SORL1 as well as experiments to determine the localization of the R953C variant withing cells. Decreased SORL1 levels are known to be pathogenic as truncation variants leading to haploinsufficiency have been definitively linked to AD [25, 26, 62]. Furthermore, human neuronal models of *SORL1* deficiency show impairments in endosomal trafficking and recycling[28, 40, 51] as do neurons from minipigs with only one functional SORL1 allele[7]. One main function of SORL1 is to sort cargo from the early endosome to either the recycling pathway (cell surface) or the retrograde pathway (TGN) in conjunction with the multi-protein complex retromer[22, 66]. The cellular localization of SORL1 and the cargo it binds depend on the specific isoform: monomer vs. dimer, mature vs. immature. For protein maturation, SORL1 transits through the Golgi and the trans-Golgi network to the endosome and to the cell surface. Here, we demonstrate that the R953C variant of SORL1 does not undergo maturation and is not shed from the cell surface.

We have recently found that two pathogenic *SORL1* missense variants associated with ADAD are located in either one of the CR-domains (Bjarnadottir et al., Manuscript in preparation) or the 3Fn-domains[35] respectively, and both display significantly impaired maturation and shedding. We have also previously observed that sSORL1 is significantly decreased in the CSF from several carriers of other established pathogenic SORL1 variants (Andersen lab, unpublished data). Furthermore, a larger screen of 70 SORL1 coding variants suggested that impaired maturation may be a common dysfunction of SORL1 mutant proteins[60].

Our study suggests that SORL1 R953C likely cannot function as a normal endosomal receptor, as it fails to enter the endosomal pathway. Instead, it is sequestered in the ER. When the receptor gets retained in the ER, it will lead to a decrease in SORL1 activity in endosomal compartments, so a direct effect of ER retention is lack-of-activity of SORL1 in the endo-lysosomal pathway. However, there could also be a gain-of-toxic-activity associated with the ER retained misfolded receptor that potentially could lead to neurotoxic ER-stress, which is suggested to occur with certain pathogenic variants in the homologous LDLR[39, 68]. Furthermore, the ER-retained SORL1 mutant protein may have additional negative impacts on total receptor activity in the endosome and thus increase the pathogenicity of the variant. In this scenario, the mutated receptor could dimerize (or even polymerize) with the wild-type receptor, thus sequestering additional wild-type SORL1 in the ER, potentially acting via a dominant-negative mechanism in diploid cells. Structural analysis indicates that this variant occurs at a critical arginine in the YWTD β-propeller domain of SORL1 that appears to be necessary for the proper folding of the domain. When compared against homologous domains in the LDLR receptor family, arginine substitutions at this position are strongly suspected to be pathogenic.

Finally, we demonstrate that this variant likely impedes SORL1 from entering the endosomal sorting pathway. SORL1 is an endosomal receptor for many proteins that are important for proper neuronal function. We and others have shown that loss of SORL1 leads to endosomal ‘traffic jams’ and mis-localization of neurotrophic receptors and glutamate receptor subunits[51]. Loss of SORL1 in the endosomal sorting pathway will likely affect multiple aspects of neuronal health and function, contributing to neurodegeneration.

Over 500 variants in *SORL1* have been identified and recent genetic studies have provided evidence as to which variants may be likely pathogenic or likely benign[25]. However, with such a large gene (encoding for more than 2200 amino acids), more variants are likely to be identified. Functional analysis of *SORL1* variants will be an important tool to classify these variants based on their cellular pathogenicity and further uncover their contribution to the development of AD.

## Acknowledgements

The authors are grateful to the family whose participation made this work possible.

Flow cytometry was performed at the FACS Core Facility, Aarhus University, Denmark.

The authors acknowledge AU Health Bioimaging Core Facility for the use of equipment and support of the imaging facility.

The authors thank the members of the University of Washington Medicine Center for Precision Diagnostics for technical support, the Geriatric Research, Education, and Clinical Center at the VA Puget Sound Health Care System, and University of Washington’s Alzheimer Disease Research Center.

We acknowledge Harald Frankowski in the Young lab for preparation of gDNA samples for SORL1 variant sequencing and all members of the Young lab for helpful discussions on this work.

