## Supplemental Fig. 1 for "A familial missense variant in the Alzheimer’s Disease gene *SORL1* impairs its maturation and endosomal sorting"

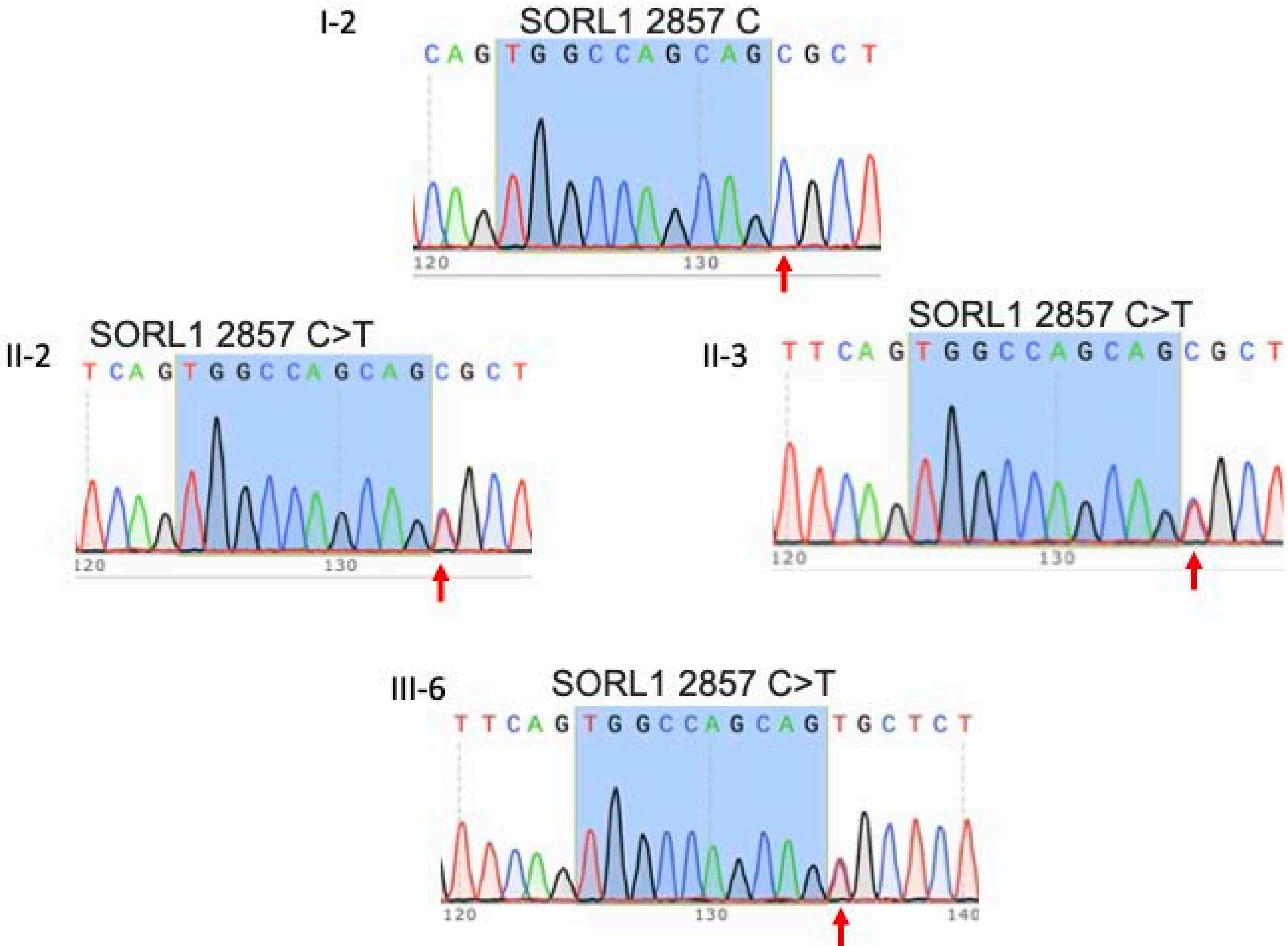
I-2
SORL1 2857 C
C A G T G G C C A G C A G C G C T
120
130
II-2
SORL1 2857 C>T
T C A G T G G C C A G C A G C G C T
120
130
II-3
SORL1 2857 C>T
T T C A G T G G C C A G C A G C G C T
120
130
III-6
SORL1 2857 C>T
T T C A G T G G C C A G C A G T G C T C T
120
130
140

**Supplemental Figure 1.** Sanger sequencing results for I-2, II-2, II-3, III-6. I-2 is C/C at *SORL1* 2857. II-2, II-3, III-6 are C/T at *SORL1* 2857.
